# Engagement rules that underpin DBL-DARC interactions for ingress of *Plasmodium knowlesi* and *Plasmodium vivax* into human erythrocytes

**DOI:** 10.1101/147769

**Authors:** Manickam Yogavel, Abhishek Jamwal, Swati Gupta, Amit Sharma

## Abstract

The molecular mechanisms by which *P. knowlesi* and *P. vivax* invade human red blood cells have long been studied. Malaria parasite erythrocytic stages comprise of repeated bursts of parasites via cyclical invasion of host RBCs using dedicated receptor-ligand interactions. A family of erythrocyte-binding proteins (EBPs) from *P. knowlesi* and *P. vivax* attach to human Duffy antigen receptor for chemokines (DARC) via their Duffy binding-like domains (*Pv*-DBL and *Pk*-DBL respectively) for invasion. Here, we provide a comprehensive overview that presents new insights on the atomic resolution interactions that underpin the binding of human DARC with *Pk*/*Pv*-DBLs. Based on extensive structural and biochemical data, we provide a novel, testable and overarching interaction model that rationalizes even contradictory pieces of evidence that have so far existed in the literature on *Pk*/*Pv*-DBL/DARC binding determinants. We address the conundrum of how parasite-encoded *Pk*/*Pv*-DBLs recognize human DARC via its two sulfated tyrosine residues. We collate evidence for two distinct DARC integration sites on *Pk*/*Pv*-DBLs that together likely engage the DARC’s sulfated extracellular domain. These analyses are important for both malaria vaccine and inhibitor development efforts that are targeted at abrogating *Pk*/*Pv*-DBL/DARC coupling as one avenue to prevent invasion of *P. vivax* into human red blood cells.

## Introduction

Engagement of specific host receptors with parasite-encoded ligands between human RBCs and malaria parasite surface proteins is a key event in invasion of merozoites into erythrocytes (Cowman *et al*., 2017). The signature Duffy-binding like domain (DBL) is present in parasite-encoded erythrocyte binding proteins (EBPs) and in protein families like *P. falciparum* EMP1s (Cowman *et al*., 2017). The latter are part of the *var* gene family where they assist in cytoadherence – these contain several copies of DBLs in each protein, whereas the *P. falciparum* EBP named EBA-175 harbors two copies (F1 and F2) of DBLs. In contrast EBPs of *P. knowlesi* and *P. vivax* contain only single DBL each. There are multiple EBPs present in the malaria parasite *P. knowelsi* (α, β and γ), and it is the *Pk*α-DBL and *Pv*-DBL that mediate DARC-dependent invasion of erythrocytes in rhesus and humans (Chao et al., 2005; Hans et al., 2005; Chitnis & Sharma 2008); a subject of discussion in this review. Other DBL variants of *P. knowelsi* EBPs: β and γ likely use alternate receptors on rhesus/human erythrocytes and may mediate invasion by Duffy antigen-independent pathways (Chitnis *et al*., 1994). The *Pk*/*Pv*-DBLs (as part of their EBPs) are present on merozoite surface and are responsible for binding to the DARC receptor on reticulocytes, and thence mediating junction formation that is vital for the parasite invasion process (Adams *et al*., 1990; Adams *et al*., 1992; Cowman *et al*., 2006) (Fig. 1). The *Pk*/*Pv*-DBLs specifically recognize DARC’s extracellular domain and its sulfated tyrosines via intermolecular interactions (Choe *et al*., 2005; de Breven *et al*., 2005; Chitnis & Sharma 2008). *Pk*/*Pv*-DBLs are organized into three sub-domains and are typified (mostly) by twelve-cysteine residues that are disulfide linked (Singh *et al*., 2003; Singh *et al*., 2006). The importance of *Pv*-DBL/DARC pairing is underscored by human genetic data where DARC negative individuals tend to be protected from *P. vivax* infection (Ryan *et al*., 2006; Cavasini *et al*., 2007; Menard *et al*., 2010; King *et al*., 2011). Contrary to the established DBL–DARC invasion pathway, there is evidence for DARC-independent invasion pathway in case of *P. vivax* infections, calling for new perspectives in assessing the utility of DARC recognizing *Pv*-DBL as a *P. vivax* vaccine candidate (Wurtz *et al*., 2011; Ntumngia *et al*., 2016).

**Figure 1:**
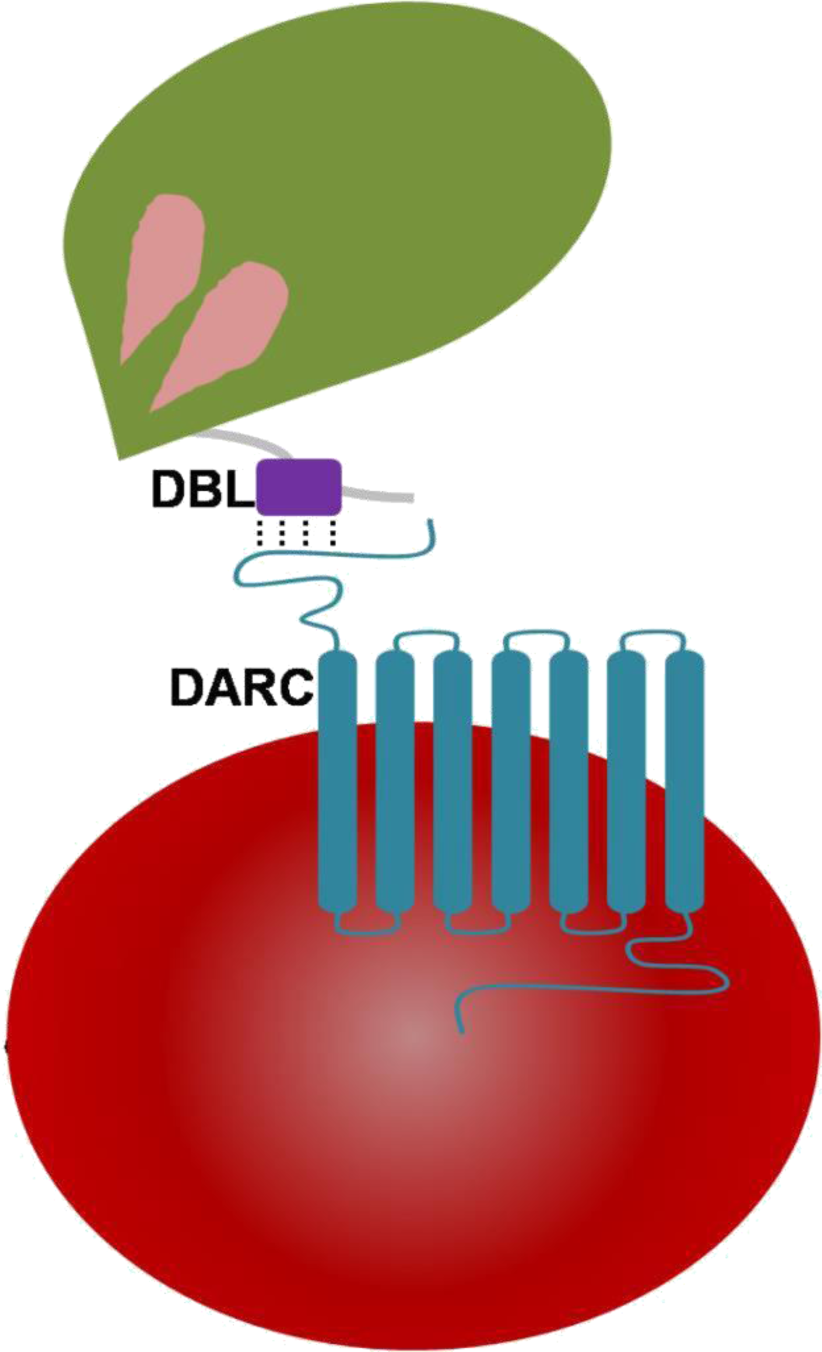
The interaction of *P. knowlesi* and *P. vivax* DBL domains with 7 transmembrane DARC on human erythrocytes. The DBLs are part of parasite-encoded membrane proteins called EBPs (erythrocyte binding proteins) that drive receptor recognition between merozoites (green) and RBCs (red). The DARC extracellular domain of residues 1-60 contains two key tyrosyl residues at positions of 30 and 41 that are post-translationally sulfated.

DARC is a seven transmembrane protein present on surface of erythrocytes and endothelial cells (Cowman *et al*., 2017)(Fig 1). It is a promiscuous cytokine/chemokine receptor involved in pro-inflammatory processes of the immune system (Rot *et al*., 2004; Cowman *et al*., 2017). DARC is also used as an entry vehicle by malaria parasites *P. knowelsi* and *P. vivax* (*Pk*/*Pv*) (Miller *et al*., 1975; Miller *et al*., 1976; Chitnis & Sharma 2008)(Fig. 1). The DARC regions spanning its soluble domain from 1-60 contain two key tyrosyl residues at positions 30 and 41, whose post-translational modifications in the form of sulfation has been reported (Choe *et al*., 2005). The binding requirements on DARC for *Pv*-DBL interaction have also been described (Choe *et al*., 2005; Tournamille *et al*., 2005). Critical *Pv*-DBL binding residues have been mapped to an N-terminal 3 region of DARC (Tournamille *et al*., 2005). Furthermore, sulfation of DARC at residues Tyr30 and Tyr41 has been strongly advocated as a key parameter for its high affinity coupling to *Pv*-DBL, where Tyr41 was shown to be the principal motif (Choe *et al*., 2005).

### Binding sites on *Pk*/*Pv*-DBLs for sulfated tyrosines of DARC

The *Pk*/*Pv*-DBLs and their three subdomains contain numerous disulfide linked cysteine residues that are largely conserved and these likely contribute to their structural integrity (Singh *et al*., 2003; Singh *et al*., 2006). Conservation of hydrophobic residues and variance in solvent exposed residues within DBLs possibly allows for a parasite-specific evolutionarily useful structural motif that can be both constant (in structural terms) and variable (in sequence). Hence DBLs seem to have evolved using the same principles as antibody structures, where the overall 3D core structures remain the same but sequence variation in exposed residues and loop regions allows for surface diversity that can thus engage with plethora of bio-molecular receptors (Singh *et al*., 2006; Gill *et al*., 2009). In efforts to understand the structural underpinnings *Pk*/*Pv*-DBL/DARC complex, the resolved crystal structure of *Pk*-DBL had suggested a region on its subdomain 2 that could accommodate the DARC’s sulfated tyrosine (Singh *et al*., 2006) and serve as its recognition site (here referred to as Site 1, Fig 2b). The *Pk*-DBL subdomain 2 presents a remarkably surface exposed region of highly conserved residues that arrange into distinct regions lying adjacent to each other – formed by positively charged residues (Lys 96, Lys 100, Arg 103 and Lys 177) and non-polars (Tyr 94, Leu 168 and Ile 175)(Singh *et al*., 2006). These two dual charge/hydrophobicity character surfaces on *Pk*-DBL were proposed to engage with DARC’s sulfated tyrosyl based on structural considerations, sequence conservation patterns, atomic properties and experimental data emanating from collation of mutagenesis experiments from two distinct groups at the time (VanBuskirk *et al*., 2004; Hans *et al*., 2005). Further, the proposed residues on *Pk*-DBL were invariant in the DARC binding DBL from *P. vivax*, hence lending support to the proposal of their essential role in acting as a recognition site for DARC’s sulfated tyrosine (VanBuskirk *et al*., 2004; Choe *et al*., 2005; Hans *et al*., 2005; Singh *et al*., 2006). These data, coupled with the observation that the sulfated Tyr41 of DARC allows strong binding to *Pk*/*Pv*-DBLs (Choe *et al*., 2005) led to a model of DARC’s docking onto *Pk*/*Pv*-DBLs via the identified site (here labeled as Site 1, Fig 2a-b). The crystal structures of *Pv*-DBLs (PDBs: 3RRC, 4NUU and 4NUV) later identified another site for DARC binding (Fig 2a, d), hereafter labeled Site 2, where based on (a) the binding of crystallization liquor phosphates (ostensibly mimicking the sulfate of DARC’s sulfated tyrosines 30 and 41), and (b) DARC peptide spanning residues 19-30, a key site for DARC binding was proposed (Batchelor *et al*., 2011; Batchelor *et al*., 2014). In these above studies, it is to be noted that the *Pk*/*Pv*-DBLs were bacterially produced, and the DARC peptide used for co-crystallization was evidently not sulfated (see PDBs 4NUU and 4NUV). The *Pv*-DBL/DARC engagement was further proposed to lead to dimerization of *Pv*-DBL (Batchelor *et al*., 2011; Batchelor *et al*., 2014). The existence of this DARC binding site (Site 2 in this review) was proposed based on the evidence of bound phosphates (PDB: 3RRC) at the (proposed) dimer interface of *Pv*-DBL (Batchelor *et al*., 2011) first, and then in a latter study of *Pv*-DBL/DARC complex, several DARC residues (numbers 19-30) were found in proximity to Site 2 (Fig 2a, d) although the bound Tyr30 was again not sulfated (Batchelor *et al*., 2011; Batchelor *et al*., 2014). The above sets of structural and biochemical data led to a conundrum on the location of the binding site for DARC’s sulfated tyrosine 41 on *Pk*/*Pv*-DBLs, and its significance in light of a previous report which had suggested that sulphation of tyrosine 41 is an essential factor for coupling of DARC with *Pk*/*Pv* DBLs (Choe *et al*., 2005). We shall return to this in a later section.

**Figure 2.**
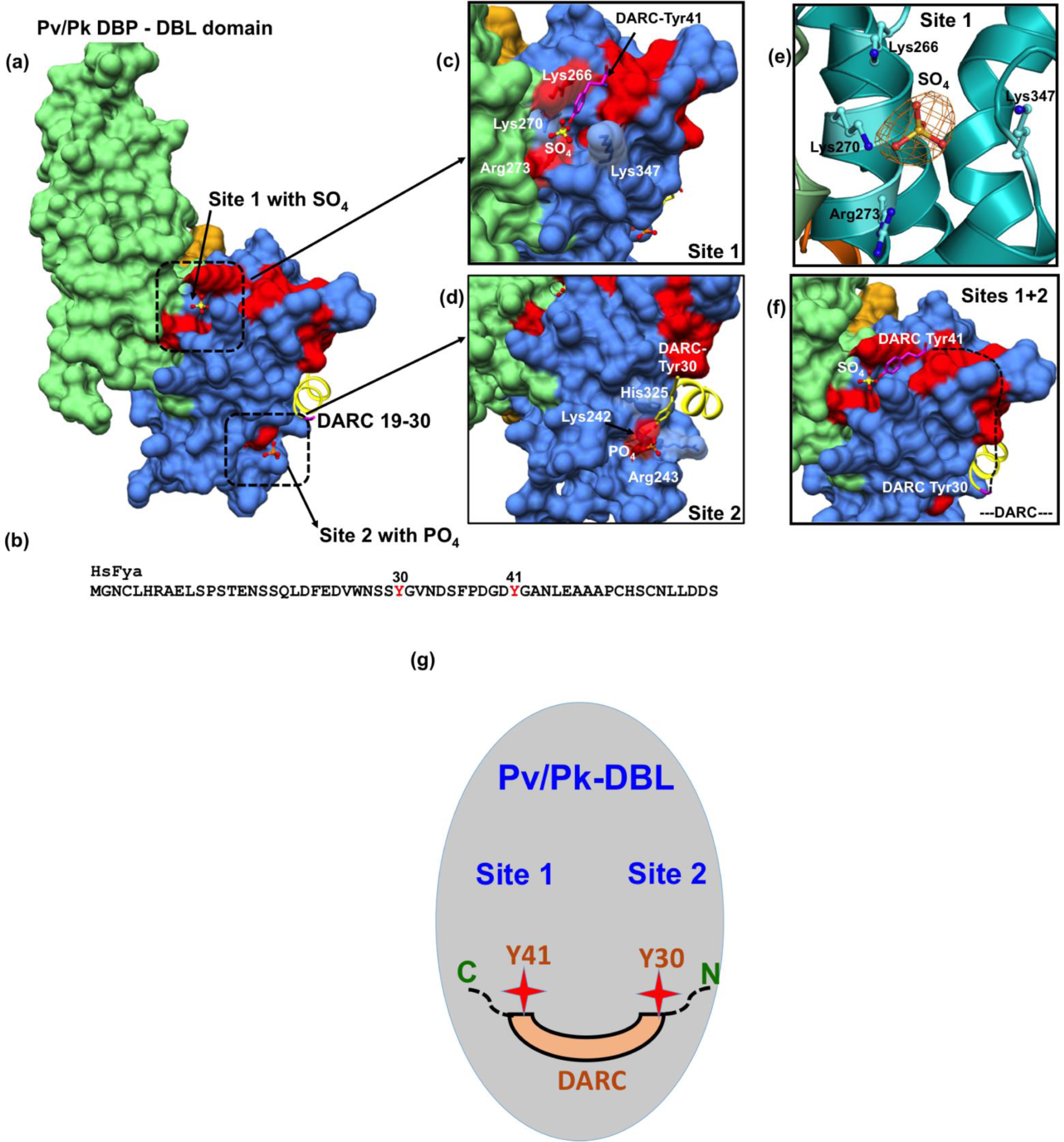
**a)** *Pk*/*Pv*-DBL subdomains 1 (orange), 2 (green) and 3 (blue) are shown as molecular surface while the N-terminal DARC peptide is shown as ribbon (yellow for duffy residues 19-30). The bound sulfate in *Pk*-DBL, phosphate in *Pv*-DBL and DARC peptide 19-30 in *Pv*-DBL are shown together. Sites 1 and 2 were identified based on mutagenesis and crystallographic data on both *Pv*/*Pk*-DBLs. The Site 2 phosphate and DARC residues 19-30 in yellow are based on crystal structures as is the sulfate found at Site 1. The underlying red patches of *Pv*-DBL residues mark the path taken by those whose mutagenesis in two different studies effects binding of *Pv*-DBL with DARC. The yellow region (19-30 of DARC) is from crystal structure of *Pv*-DBL/DARC complex while the purple DARC represents modeling such that sulphated-Tyr41 is proximal to the sulfate observed in *Pk*-DBL structure. The DARC peptide can easily traverse from Tyr30 to Tyr41 as it docks on to Sites 2 and 1 respectively. **b)** Sequence of human DARC with tyrosyl residues 30 and 41 shown in red as these are proposed to be important for both chemokine and *Pv*-DBL binding. Both tyrosines 30 and 41 are sulfated in human DARC and it has been proposed that tyrosine 41 sulfation is important for high affinity *Pk*/*Pv*-DBL/DARC engagement. **c)** Zoomed view of Site 1 on *Pv*/*Pk*-DBLs with sulfate interacting residues that are identical between *Pk*/*Pv*-DBLs. Note that the corresponding residue numbers are for *Pk*-DBL (PDB: 5X6N). The DARC-tyrosine 41 is modeled to show its proximity to the sulfate position found in crystal structure of *Pk*-DBL (PDB: 5X6N). The underlying red patches of *Pv*-DBL residues mark the path taken by those whose mutagenesis in two different studies effects binding of *Pv*-DBL with DARC. **d)** Zoomed view of Site 2 on *Pv*-DBL modeled from superposition of two *Pv*-DBL structures (PDB IDs: 3RRC and 4NUV) with bound phosphate and unsulfated DARC peptide (residues 1930). Note that the corresponding residue numbers are for *Pv*-DBL (PDB: 3RRC). The underlying red patches of *Pv*-DBL residues mark the path taken by those whose mutagenesis in two different studies effects binding of *Pv*-DBL with DARC. **e)** The F_0_-F_c_ electron density (green) and SA-OMIT (orange) map contoured at 3 sigma level for bound SO_4_ in *Pk*-DBL (PDB: 5X6N) at Site 1. Note that the corresponding residue numbers are for *Pk*-DBL (PDB: 5X6N). **f)** Zoomed view of *Pv*-DBL where modeling has resulted in superposition of bound DARC peptide (19-30) and bound sulfate (from *Pk*-DBL). The dashed line shows the proposed trajectory of DARC peptide that it may take once its Tyr30 is bound to Site 2 and its Tyr41 is bound to Site 1. Again, note that many residues that are critical (colored red) for DARC binding, as revealed by two separate mutagenesis studies, fall along the path from Site 2 to Site 1 resulting in a wide DARC binding footprint. **g)** Proposed model on the modes of interaction between *Pv*/*Pk*-DBLs and sulfated DARC. We suggest that the DARC peptide, via its sulfated tyrosines 30 and 41, engages at two subsites (Sites 2 and 1 respectively) on *Pv*/*Pk*-DBLs. Sites 1 and 2 are reasonably spaced for binding of intervening DARC residues between tyrosines 30 and 41. This testable model is a dual-site mode of DARC’s engagement with *Pv*/*Pk*-DBLs and can be experimentally assessed now.

It has been also proposed that *Pv*-DBL engagement with DARC extracellular domain drives dimerization of *Pv*-DBLs (PDB IDs: 3RRC, 4NUU and 4NUV) (Batchelor *et al*., 2011; Batchelor *et al*., 2014). Extensive analysis using PISA (http://www.ebi.ac.uk/pdbe/pisa/) and EPPIC (http://www.eppic-web.org/), two softwares that analyze the oligomeric states of biomolecular structures deposited in the public database PDB (Krissinel *et al*., 2007; Duarte *et al*., 2012; Baskaran *et al*., 2014), does not support dimerization of *Pv*-DBL when bound to phosphate or to the duffy peptide spanning residues 19-30 (we strongly encourage readers to assess this using online PISA and EPPIC softwares and the PDB: 3RRC, 4NUU and 4NUV).However, we do not preclude the oligomerization of *Pv*-DBLs when bound to DARC *in vivo*, and/or in the context of full-length proteins of each when they interact on their respective cell surfaces. Yet from available structural data, it is evident that the *Pv*/*Pk*-DBL monomer contains all vital structural elements required for engagement with two sites on sulfated DARC. Indications of dimeric states gathered from in solution studies like SAXS, ITC and analytical ultracentrifugation do not necessarily reveal the exact binding interfaces in a biomolecular complex. The structural resolution of *Pk*/*Pv*-EBPs bound to fully sulfated DARC will eventually reveal the modes of complex formation between these receptor ligand pairs.

### A new overarching model for coupling of *Pk*/*Pv*-DBLs with DARC

The *Pv*-DBL (PDB: 3RRC) has two bound phosphates near/at Site 2, where later DARC peptide residues (19-30) were also found (PDBs: 4NUU, 4NUV). During process of routine structure re-refinement of deposited crystal structures, as recommended (Read et al., 2011) due to the more robust convergence routines and higher quality of resulting stereochemical models for proteins, we observed a bound sulfate at Site 1 in the *Pk*-DBL crystal (Fig 2a, c). The bound sulfate site overlaps with the earlier proposed sulfa-tyrosine recognition region (Singh *et al*., 2006), nestled by the highly conserved positively charged residues most of which were earlier mapped based on mutagenesis screens (VanBuskirk *et al*., 2004; Choe *et al*., 2005; Hans *et al*., 2005; Singh *et al*., 2006). Indeed, both *Pv*/*Pk*-DBLs have other phosphate/sulfate binding sites in their crystal structures but we have taken into account only those that are proximal to the regions where mutagenesis and biochemical data provide strong evidence for DARC engagement (VanBuskirk *et al*., 2004; Hans *et al*., 2005; Sampath *et al*., 2013). The *Pk*-DBL protein migrates as a monomer on gel filtration column in the absence of detergent, and in the crystal it is also an unambiguous monomer – even when bound to a SO_4_ in Site 1 (see PDB ID: 5X6N and assess via PISA/EPPC). The detergent binding site in 5X6N is in any case far from Sites 1 and 2 (see PDB ID: 5X6N). The presence of bound sulfate at Site 1 in *Pk*-DBL is clearly a striking observation, and it immediately opens the possibility of reassessing and reinterpreting the engagement rules for DARC recognition and binding by *Pk*/*Pv*-DBLs.

Therefore, based on (a) *Pk*-DBL-sulfate complex (Site 1, Fig. 2a, c, e), (b) *Pv*-DBL-phosphate complex (Site 2, Fig. 2a, d), (c) *Pv*-DBL-DARC peptide complex (Site 2, Fig. 2b, d) and available mutagenesis and biochemical data on *Pv*-DBL/DARC interactions (VanBuskirk *et al*., 2004; Hans *et al*., 2005; Sampath *et al*., 2013), we are able to propose a simple, novel, testable and fully rationalized resolution to the conundrum of *Pk*/*Pv*-DBL/DARC coupling (Fig. 2f, g). We envisage that the DARC peptide 1-60 may dock on to *Pk*/*Pv*-DBLs via its sulphated tyrosine 41 on Site 1 (Fig. 2a, f, g; PDB ID: 5X6N) and on to Site 2 via the DARC peptide containing residues 19 to Tyr 30 (Fig 2a, e, f; PDB ID: 4NUV). Structural, molecular size and stereo-chemical considerations allow the DARC peptide from residue 19-41 to traverse from Site 2 (where tyrosine 30 is) to Site 1 (where tyrosine 41 is) on *Pk*/*Pv*-DBLs. This dual site DARC hooking on to *Pk*/*Pv*-DBLs offers a more complete binding footprint of DARC (Fig 2f, g). Indeed, the molecular size of a sulfated tyrosine (from its C-alpha to sulfate) is ~14Å. Quick modeling of the atomic space between *Pk*-DBL’s Y94 (Site 1 residue, as proposed Singh *et al*., 2006) and the bound sulfate shows that a sulfated tyrosine will fit snugly into it. Further, the side chains in *Pk*/*Pv*-DBLs that constitute Site 1 are identical (4/4), and mostly (2/3) conserved for Site 2 (Fig 2 c,d). Invariance in Site 1 residues of *Pk*/*Pv*-DBLs: Lys96, Lys100, Arg103 and Lys 177 (*Pk*-DBL numbering, as in Singh *et al*., 2006) that nestle (i.e charge complementarity) the bound sulphate moiety (Fig 2e), and its potential as a sulfa-tyrosine recognition site based on earlier mutagenesis experiments together provide very strong support for the significance of Site 1 in *Pk*/*Pv*-DBL/DARC interactions. These analyses hence propose that both Sites 1 and 2 are part of the overall binding footprint of DARC on *Pv*/*Pk*-DBLs, and that Sites 1 and 2 likely represent the regions of engagement where sulfated Tyr41 and Tyr30 (respectively) are recognized by the *Pv*/*Pk*-DBLs (Fig 2f). The intervening regions of *Pv*/*Pk*-DBL and DARC will also likely make contact with one another, and therefore the mutagenesis footprint on DBLs will be larger than that alone for Sites 1 and 2 – as indeed seems to be the case, and shall be discussed below (VanBuskirk *et al*., 2004; Hans *et al*., 2005).

Earlier, single amino acid mutagenesis data had suggested over ˜two dozen mutations that could effect binding of DARC to *Pv*-DBL (VanBuskirk *et al*., 2004; Hans *et al*., 2005). Based on comprehensive collation of structural and experimental data, we now propose a model that shows that majority of the 25 mutations (VanBuskirk *et al*., 2004; Hans *et al*., 2005) that diminish *Pv*-DBL/DARC engagement fall in a predicted trajectory that DARC peptide may take when traversing from Site 1 from Site 2 (Figure 2f). Two reports had earlier identified several sets of residues on *Pv*-DBL that altered binding to DARC – several of these residues line and nestle the sulfate in *Pk*-DBL at Site 1, besides being proximal to Site 2 (VanBuskirk *et al*., 2004; Hans *et al*., 2005). Further support for our model comes from a glycan masking study that was focused on *Pv*-DBL/DARC wherein it was suggested that addition of an N-glycan site near the proposed Site 1, but not in other regions of *Pv*-DBL, including the proposed dimer interface site (Site 2), abrogated DARC binding (Sampath *et al*., 2013). These data additionally support our proposed model of twin engagement of DARC via its sulfated tyrosine 41 at Site 1 and with DARC residues 19-30 at Site 2: comprehensive mapping of mutagenesis data support our idea that DARC residues from 31 to 40 traverse from Site 2 towards Site 1 along the path where many mutants map (Fig. 2f).

It needs to be emphasized again that our proposed model is not discounting the importance of DARC (residues 19-30) identified earlier (Tournamille *et al*., 2005; Batchelor *et al*., 2014). Our new suggested structural framework of *Pk*/*Pv*-DBL/DARC interactions here supports, collates and integrates many studies (VanBuskirk *et al*., 2004; Choe *et al*., 2005; Hans *et al*., 2005; Singh *et al*., 2006; Batchelor *et al*., 2011; Batchelor *et al*., 2014). Our model is consistent with most mutagenesis and crystallography work published so far, and provides a more comprehensive explanation for DARC’s binding via its sulfated Tyr30 and Tyr41 to two binding sites (Sites 1 and 2) each on *Pk*/*Pv*-DBLs (Fig 2a-g). The proposed architecture of the complex sheds new light on DBL-DARC interactions and our overview suggests avenues for experimental validation. Our proposal also offers scope to make new sets of mutants that can directly address these hypotheses using the available 3D structures.

Finally, the indicated structural framework of *Pk*/*Pv*-DBL/DARC significantly resolves the puzzle of recognition site (s) that underpin this complex. Our analyses integrate new evidence for two distinct DARC binding sites on *P. knowlesi* and *P. vivax* DBLs (called Sites 1 and 2, in the chronological order they were described in the literature) that together likely engage the extracellular domain of DARC via sulfated tyrosines 41 and 30 respectively (fig 2g). Our analyses shall be important for critical assessment of both malaria vaccine and inhibitor development efforts that are targeted at abrogating *Pk*/*Pv*-DBL/DARC interactions as a possible avenue to prevent invasion of *Pk*/*Pv* malaria parasites into human red blood cells.

